# Heart rate variability as an indicator of autonomic nervous system disturbance in tetanus

**DOI:** 10.1101/793497

**Authors:** Ha Thi Hai Duong, Girmaw Abebe Tadesse, Phung Tran Huy Nhat, Nguyen Van Hao, John Prince, Tran Duc Duong, Trịnh Trung Kien, Le Van Tan, Chris Pugh, Huynh Thi Loan, Nguyen Van Vinh Chau, Yen Lam Minh, Tingting Zhu, David Clifton, Louise Thwaites

**Affiliations:** Hospital for Tropical Diseases, Ho Chi Minh City, Viet Nam; Institute of Biomedical Engineering, University of Oxford, UK; Oxford University Clinical Research Unit, Ho Chi Minh City, Viet Nam; University of Medicine and Pharmacy, Ho Chi Minh City, Viet Nam; Nuffield Department of Medicine, University of Oxford, UK; Centre for Tropical Medicine and Global Health, University of Oxford, UK

## Abstract

Autonomic nervous system dysfunction (ANSD) is a significant cause of mortality in tetanus. Currently diagnosis relies on non-specific clinical signs. Heart rate variability (HRV) may indicate underlying autonomic nervous system activity and represents a potentially valuable non-invasive tool for ANSD diagnosis in tetanus. HRV was measured from 3 5-minute ECG recordings during a 24-hour period in a cohort patients with severe tetanus, all receiving mechanical ventilation. HRV measurements from all subjects - 5 with ANSD (Ablett Grade 4) and 4 patients without ANSD (Ablett Grade 3) - showed HRV was lower than reported ranges for healthy individuals. Comparing different severities of tetanus, raw data for both time and frequency measurements of HRV were reduced in those with ANSD compared to those without. Differences were statistically significant in all except root mean square standard deviation RMSSD (p=0.07) indicating HRV may be a valuable tool in ANSD diagnosis.

## Introduction

Tetanus is a severe disease characterized by toxin-mediated disinhibition of autonomic and motor nervous systems. Motor neuron disinhibition causes characteristic muscle spasms, whereas autonomic nervous system disinhibition results in fluctuating blood pressure, tachycardia and pyrexia. When mechanical ventilation is available, spasms can be controlled but ANSD remains a principal cause of mortality ^1,2^. Robust methods of detecting ANSD suitable for implementation in resource-limited settings where most tetanus occurs would allow earlier intervention and may improve outcome. Diagnosis is currently based on non-specific clinical signs of pyrexia, sweating and increased or fluctuating heart rate and blood pressure ^3^. Other methods include 24-hour collections of urinary catecholamines, but this has low specificity, unsuitable for routine use ^4^.

In health, heart rate is carefully controlled by the autonomic nervous system. Alterations in parasympathetic and sympathetic nervous system activity result in beat-to-beat heart rate variation and hence this variation (heart rate variability) reflects autonomic nervous system activity. Heart rate variability (HRV) is altered in pathological states such as ischaemic heart disease and reduced variability is predictive of worse outcomes ^5^. Standardized measures of HRV can be calculated from electrocardiogram (ECG) R-R intervals and consensus guidelines on appropriate indicators are available ^5^. Time domain variables are calculated directly from R-R intervals (termed normal-to-normal intervals), for example standard deviation. Frequency domain variables are generated from ECG spectral analysis, usually following Fast Fourier Transformation ^5^. By observing changes in these components after administering autonomic nervous system antagonists, relative contributions of parasympathetic and sympathetic nervous systems has been inferred. Whilst total power of the spectrum represents the general level of autonomic activation, low frequency activity (<0.15Hz) is mainly due to baroreceptor reflex modulation and related to both vagal and sympathetic influence, whereas high frequency activity is mainly aligned with vagal activity. Low:high frequency ratio is accepted to indicate balance between both systems; however, this interpretation fails to take account of effects such as different temporal patterns of sympathetic and parasympathetic components and cardiac pacemaker sensitivity.

HRV changes in tetanus are largely unknown. Sykora *et al* analysed baroreflex sensitivity and time domain variables in an 87 year-old woman with tetanus and reported decreased baroreceptor sensitivity compared to a control of similar age, however the patient, but not control, received mechanical ventilation and a beta blocker both of which can influence sensitivity ^6^. Goto *et al* reported reduced frequency-domain variables in an 11 year-old child, however this recording was taken following a cardiac arrest and on the 122^nd^ day of hospitalization where clinical recovery from tetanus is normally expected ^7^.

Nevertheless, ANSD diagnosis and prognostication through HRV remains an attractive prospect due to its non-invasive nature. Hitherto, required monitoring equipment was rarely available in settings where most tetanus occurs, but growing availability of low-cost sensors means measurement is increasingly feasible in low-resource settings ^8^. In this study, we aim to investigate the relationship of HRV and ANSD in patients with severe tetanus, providing proof-of-principal that such monitoring may be valuable.

## Methods

The study was conducted in the Intensive Care Unit at the Hospital for Tropical Diseases, Ho Chi Minh City between October 2016 and January 2017 and was approved by the Ethical Committee of the Hospital for Tropical Diseases. Written informed consent was given by all participants or representatives before enrolment.

Adults with severe tetanus (Ablett Grade 3 or 4), diagnosed according to the Hospital for Tropical Disease guidelines ^9,10^ and receiving mechanical ventilation were recruited to the study. Recruitment was pragmatic and depended on availability of suitable monitors. Ablett Grade 3 was defined as ‘severe spasms interfering with respiration’ and Grade 4 as grade 3 but with ANSD ^10^. ANSD was diagnosed clinically by the attending physician but required the presence of at least 3 of the following within 12 hours: heart rate > 100 bpm, systolic blood pressure > 140 mmHg, blood pressure fluctuation with minimum mean arterial pressure < 60 mmHg, temperature > 38°C without evidence of intercurrent infections.

Tetanus management followed a standard protocol previously described ^11^, consisting of antibiotics, spasm control using benzodiazepines and pipecuronium. ANSD was managed principally with magnesium sulphate.

ECG data were collected from bedside monitors (Datex, Datex Ohmeda Inc, USA) in supine undisturbed patients using VSCapture Software ^12^. ECG, physiological and clinical data were collected over a 24-hour period. HRV features were extracted from noise-free 5-minute recordings at 6am, 12 noon and 6pm to prevent bias from HRV diurnal variation^13^. Time domain variables measured were square root of the mean squared differences of successive normal-to-normal intervals (RMSSD) and standard deviation of all normal-to-normal intervals (SDNN). Frequency domain variables were total power, high frequency power (0.15-0.4 Hz) low frequency power (0.05-0.15Hz), low frequency normalized units, high frequency normalized units and low: high frequency ratio. Statistical analyses were performed using R statistical software version 3.5.1 (R corporation). Data are presented as mean (SD). Heart rate variability was compared using analysis of multiple variance (ANOVA). A p value < 0.05 was considered statistically significant.

## Results

Five patients with Ablett Grade 4 and 5 patients with Ablett Grade 3 tetanus were recruited to the study. Data from one patient with Grade 3 tetanus was too noisy for analysis and therefore excluded. Clinical characteristics of the remaining 9 patients are given in Table 1. Of these, 8/9 had 3 high-quality noise-free 5-minute segments at the chosen time point. One patient with Ablett Grade 4 had only 2 suitable 5-minute segments at 12 noon and 6pm.

**Table 1.** Clinical data of patients. Figures given are mean (SD), except males, mechanical ventilation and mortality which are n (%).

|  | **Ablett\* Grade 3 n=4** | **Ablett\* Grade 4 (ANSD) n=5** |
| --- | --- | --- |
| **Age (years)** | 51.25 (20.66) | 58 (7.68) |
| **Males** | 4 (100) | 4 (80) |
| **Tetanus Severity Score** | 5.8 (6.5) | 4.8 (4.09) |
| **Mechanical ventilation** | 4 (100) | 5 (100) |
| **In-hospital mortality (%)** | 0 (0) | 0 (0) |
| **Maximum heart rate in 24-hour study period (bpm)** | 152.8 (22.34) | 210.2 (29.26) |
| **Minimum heart rate in 24-hour study period (bpm)** | 59.00 (6.21) | 48.20 (10.08) |
| **Mean heart rate in 24-hours study period (bpm)** | 94.50 (10.25) | 90.00 (20.41) |
| **Mean systolic blood pressure in 24-hour study period (mmHg)** | 127.60 (2.27) | 159.90 (21.06) |
| **Maximum systolic blood pressure in 24 hours study period (mm Hg)** | 137.50 (9.57) | 217.40 (26.49) |
| **Minimum systolic blood pressure in 24 hours study period (mm Hg)** | 115.00 (5.77) | 107.40 (27.24) |
| **Total dose of diazepam during 24-hour study period (mg)** | 30.00 (60) | 48.00 (65.72) |
| **Total dose of midazolam during 24-hour study period (mg)** | 120.00 (80.80) | 86.40 (84.17) |
| **Total dose of magnesium sulphate during 24-hour study period (mg)** | 0 (0) | 43.20 (26.29) |
| **Total dose of pipecuronium during 24-hour study period (mg)** | 38.40 (7.83) | 30.72 (4.29) |

\* Ablett Grade 3: severe tetanus with spasms compromising respiration. Ablett Grade 4 is as Grade 3 but with additional signs of autonomic nervous system dysfunction.

HRV data are presented in Figure 1 and Table 2. All HRV measurements were very low compared to reported ranges for healthy individuals, with low:high frequency ratios significantly greater ^5^. Comparing different severities of tetanus, both time (RMSSD, SDNN) and frequency (low frequency, high frequency, low frequency normalized units, low:high frequency and total power) variables were reduced in those with ANSD (Ablett Grade 4) compared to those without. Differences were statistically significant in all except RMSSD (p=0.07). Only high frequency normalized units showed no difference between groups (p=0.9).

**Table 2:** Heart rate variability variables. Variables presented are mean (SD)

|  | **Ablett Grade 3, n=12** | **Ablett Grade 4 (ANSD), n=14** | **p** |
| --- | --- | --- | --- |
| **Low frequency**<br>**(normalized units)** | 62.54 | 27.29 | 0.001 |
| **High frequency**<br>**(normalized units)** | 21.26 | 22.03 | 0.92 |
| **Low:high frequency ratio** | 8.59 | 2.75 | 0.18 |
| **High frequency (ms<sup>2</sup>)** | 43.28 | 7.32 | 0.005 |
| **Low frequency (ms<sup>2</sup>)** | 227.57 | 21.01 | 0.03 |
| **Total power (ms<sup>2</sup>)** | 609.77 | 150.81 | 0.01 |
| **RMSSD (ms)** | 15.33 | 6.95 | 0.08 |
| **SDNN (ms)** | 28.82 | 15.67 | 0.002 |

**Figure 1:**
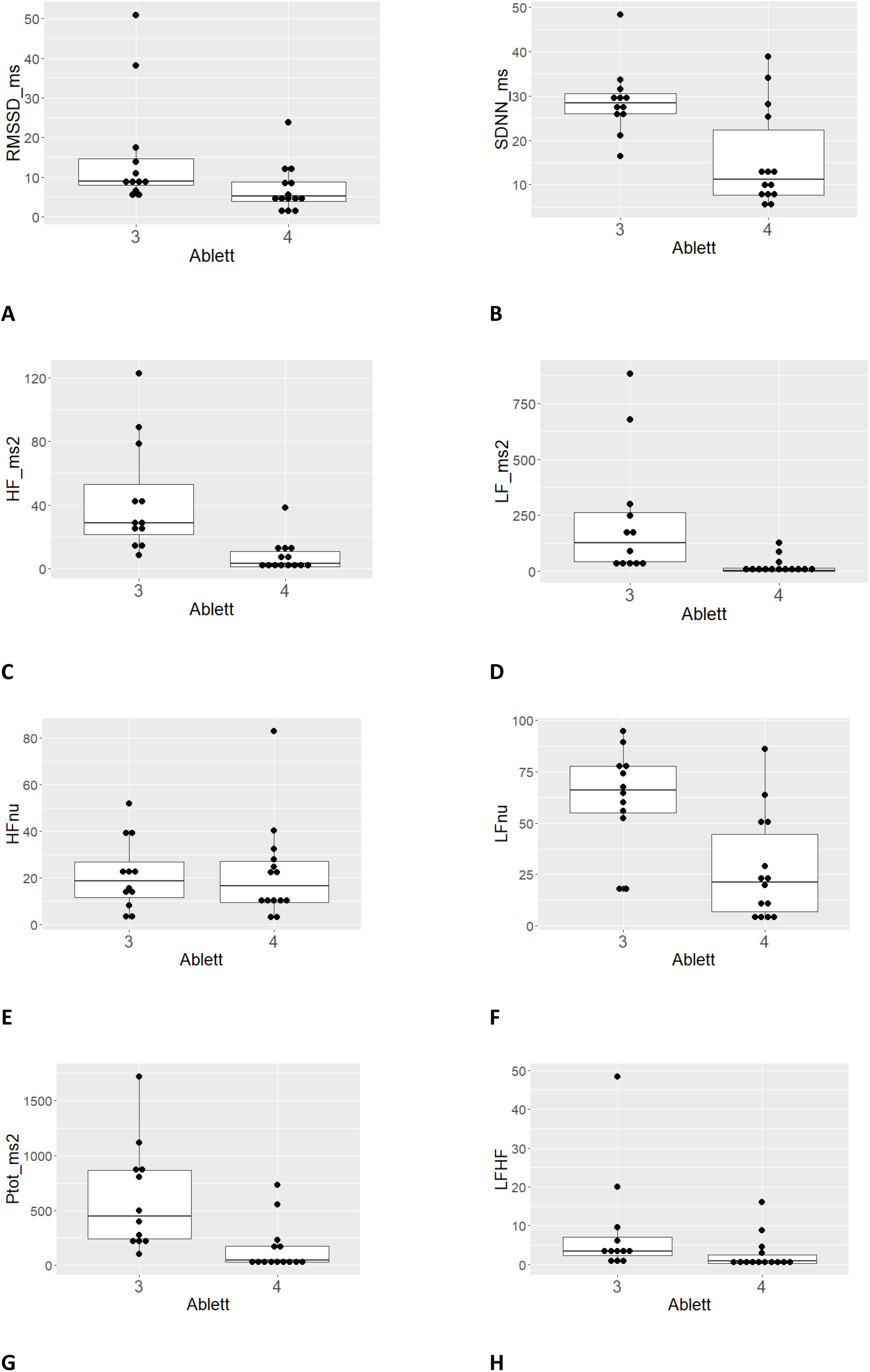
Heart rate variability variables in patients according to Ablett Grade. Panel A, square root of the mean squared differences of successive normal-normal (NN) intervals (RMSSD); Panel B, standard deviation of all NN interval (SDNN), Panel C: Power in high frequency range; Panel D, power in low frequency range; Panel E, high frequency power in normalized units; Panel F, low frequency power in normalized units, Panel G, total spectral power and H, low: high frequency ratio.

## Discussion

We present, to our knowledge, the first HRV measurements in a series of patients with tetanus. Our data show a consistent reduction in time and frequency domain variables compared to values reported in healthy subjects. These are particularly reduced in those with clinical signs of ANSD. This is consistent with HRV reported in other pathological states with high levels of sympathetic activation and with existing understanding of ANSD in tetanus.

Sympathetic activation in tetanus is associated with increased circulating catecholamines, which are increased in proportion to disease severity ^4^. These may exert direct effects on the heart and vasculature and indirect effects through reflex reduction in vagal tone. The observed reduction in HRV variables in those with ANSD is consistent with sympathetic nervous system activation. Whilst the reduction in high frequency power, suggesting a reduction in vagal tone is expected, we also observed a reduction in low frequency power, indicative of both sympathetic and parasympathetic activation. In cases of sympathetic activation, heart rate increases and total power is reduced and, as a result, the low frequency component may actually fall ^14^. Similarly at high levels of sympathetic stimulation a ‘ceiling effect’ may occur at the sinoatrial node when further response cannot occur ^14^.

A significant limitation to interpretation of our data is that our patients were all receiving sedative drugs which may influence heart rate variability. Although, sedation is not reported to affect HRV in critically ill patients ^15^ and subjects in both groups received similar sedative doses, it is possible that drugs were titrated against clinical effect. Magnesium sulphate was used almost exclusively in those with ANSD. Although we have previously shown its use in tetanus is associated with a reduction in urinary catecholamine excretion, limited data in myocardial infarction suggest that it has limited effect on HRV ^16,17^.

A further limitation is that we used 5-minute recordings to measure time domain variables. Guidelines recommend that these should be measured from 24-hour recordings. Nevertheless our values are lower than reported 5-minute ‘normal’ values and our primary comparison was between severity groups ^5^.

Heart rate control in tetanus is undoubtedly complex and influenced by many factors not measured in this study. As such, we aimed only to demonstrate that variability is related to disease severity and that alterations in HRV may aid ANSD diagnosis in patients with tetanus. Nevertheless HRV could potentially be a more sensitive and specific way of identifying those with ANSD. The low HRV observed even in patients with Grade 3 tetanus may represent ‘subclinical’ ANSD where intervention may be beneficial. Furthermore, HRV changes may be early predictors of subsequent ANSD and enable earlier intervention.

This paper has focused on using established HRV measures, however it is likely these are relatively blunt tools with which to decipher the complex underlying physiological mechanisms and rely on high-quality signals difficult to obtain in critically ill populations in resource-limited settings. As newer innovative methods for analysing data develop, for example artificial intelligence, more sensitive ways of analysis could be developed and providing better insight into control mechanisms and disease pathophysiology.

## Acknowledgments

We thank all staff in adult AICU at the Hospital Diseases and the biostatistics group at Oxford University Clinical Research Unit for statistical advice.

